# RENET2: High-Performance Full-text Gene-Disease Relation Extraction with Iterative Training Data Expansion

**DOI:** 10.1101/2021.03.18.436005

**Authors:** Junhao Su, Ye Wu, Hing-Fung Ting, Tak-Wah Lam, Ruibang Luo

**Affiliations:** Department of Computer Science, The University of Hong Kong, Hong Kong, China

## Abstract

**Background:** Relation extraction is a fundamental task for extracting gene-disease associations from biomedical text. Existing tools have limited capacity, as they can extract gene-disease associations only from single sentences or abstract texts.

**Results:** In this work, we propose RENET2, a deep learning-based relation extraction method, which implements section filtering and ambiguous relations modeling to extract gene-disease associations from full-text articles. We designed a novel iterative training data expansion strategy to build an annotated full-text dataset to resolve the scarcity of labels on full-text articles. In our experiments, RENET2 achieved an F1-score of 72.13% for extracting gene-disease associations from an annotated full-text dataset, which was 27.22%, 30.30% and 29.24% higher than the best existing tools BeFree, DTMiner and BioBERT, respectively. We applied RENET2 to (1) ~1.89M full-text articles from PMC and found ~3.72M gene-disease associations; and (2) the LitCovid articles set and ranked the top 15 proteins associated with COVID-19, supported by recent articles.

**Conclusion:** RENET2 is an efficient and accurate method for full-text gene-disease association extraction. The source-code, manually curated abstract/full-text training data, and results of RENET2 are available at https://github.com/sujunhao/RENET2.

## Introduction

The association between genes and diseases is essential for developing clinical diagnoses, therapeutic treatments, and public health systems for diseases [1]. However, the research on gene-disease associations is locked in an enormous volume of biomedical literature. PubMed Central (PMC) [2], a free full-text archive of the biomedical literature, had over 6.6 million articles in 2020. There is a pressing need for accurate and efficient tools to automatically extract gene-disease associations from the literature to improve access to information and support biomedical research [3].

The Relation Extraction (RE) task is critical for extracting gene-disease associations from the literature [4]. The task is to determine whether there is an association between a gene-disease pair from a given text. Relation extraction is more challenging than the task of finding named entities from texts, namely Named Entity Recognition (NER [5, 6]), as it has to incorporate the information from complete sentences (sentence-based) or complete articles (document-based). A wide range of methods, such as BeFree [7], DTMiner [8], and LHGDN [9], employ a sentence-based approach to extracting gene-disease relations. These methods utilize different linguistic and co-occurrence features with machine-learning methods to identify gene-disease relations within each sentence. For example, BeFree [7] applies a shallow linguistic kernel, which uses both a local (orthographic and shallow linguistic features) and global context (trigrams and sparse bigrams) to extract relations from a single sentence. DTMiner [8] improves BeFree by adding a co-occurrence-based ranking module to estimate how closely the pairs are related. However, sentence-based methods can extract relations only within a sentence. In an article, information about gene-disease associations is often spread over multiple sentences. To extract gene-disease associations supported by an article, we need document-based relation extraction methods to understand the context of the whole article.

Existing document-level relation extraction methods are designed mainly for abstract texts. BioBERT [10] is a comprehensive approach, which applies BERT [11], an attention-based language representation model [12], on biomedical text mining tasks, including Named Entity Recognition (NER), Relation Extraction (RE), and Question Answering (QA) [13]. BioBERT can extract gene-disease associations from biomedical text by performing classification on both sentences and abstracts. However, because of its attention mechanism, BioBERT constrains its maximum input length to 512 and restricts its application to complete articles. The predecessor to this work, called RENET [14], uses a Convolutional Neural Network (CNN) and a Recurrent Neural Network (RNN) to learn the document representation of gene-disease relations. RENET not only captures the relationship between genes and diseases within a sentence, but also models the interaction of different sentences to understand the context of the whole article. It achieves state-of-the-art performance in gene-disease relation extraction from abstracts. However, RENET uses classification, which is similar to BioBERT, to model the relation extraction problem. This mechanism is not flexible enough to handle more complex relation types, such as ambiguous relations. Moreover, it is designed for abstract texts, not optimized for full-text articles. The existing methods still pose significant challenges to full-text relation extraction [15].

Full-text articles contain much more information than abstract texts. We estimate that a full-text article has an average of 4,535 tokens, about 17 times that of an abstract. The average number of gene-disease pairs in a full-text article is estimated to be 197, more than 60 times that of an abstract. This rich content makes gene-disease relation extraction from full text more challenging than from abstracts. The length of full-text articles exceeds the capacity of BioBERT and other BERT-based methods. It reduces the information density of the text, resulting in low prediction accuracy. The long length and large number of gene-disease pairs also make manual curation of a full-text dataset labor-intensive and time-consuming. As a result, there is no publicly available full-text level labeled data for development and evaluation.

In this paper, we propose RENET2, an accurate and efficient full-text gene-disease relation extraction method with an iterative training data expansion strategy, as shown in Figure 1. In RENET2, we introduced “Ambiguous association”, a new relation type to reduce human effort in labeling gene disease associations, and used a regression-based deep-learning approach to model Ambiguous associations. We applied section filtering, a novel data-enhancement technique, to reduce the noisy content and improve the information density of the input data for gene-disease relation extraction. We designed a training data expansion strategy, which performs model training, prediction and efficient manual curation iteratively to generate ample high-quality full-text training samples.

**Figure 1.**
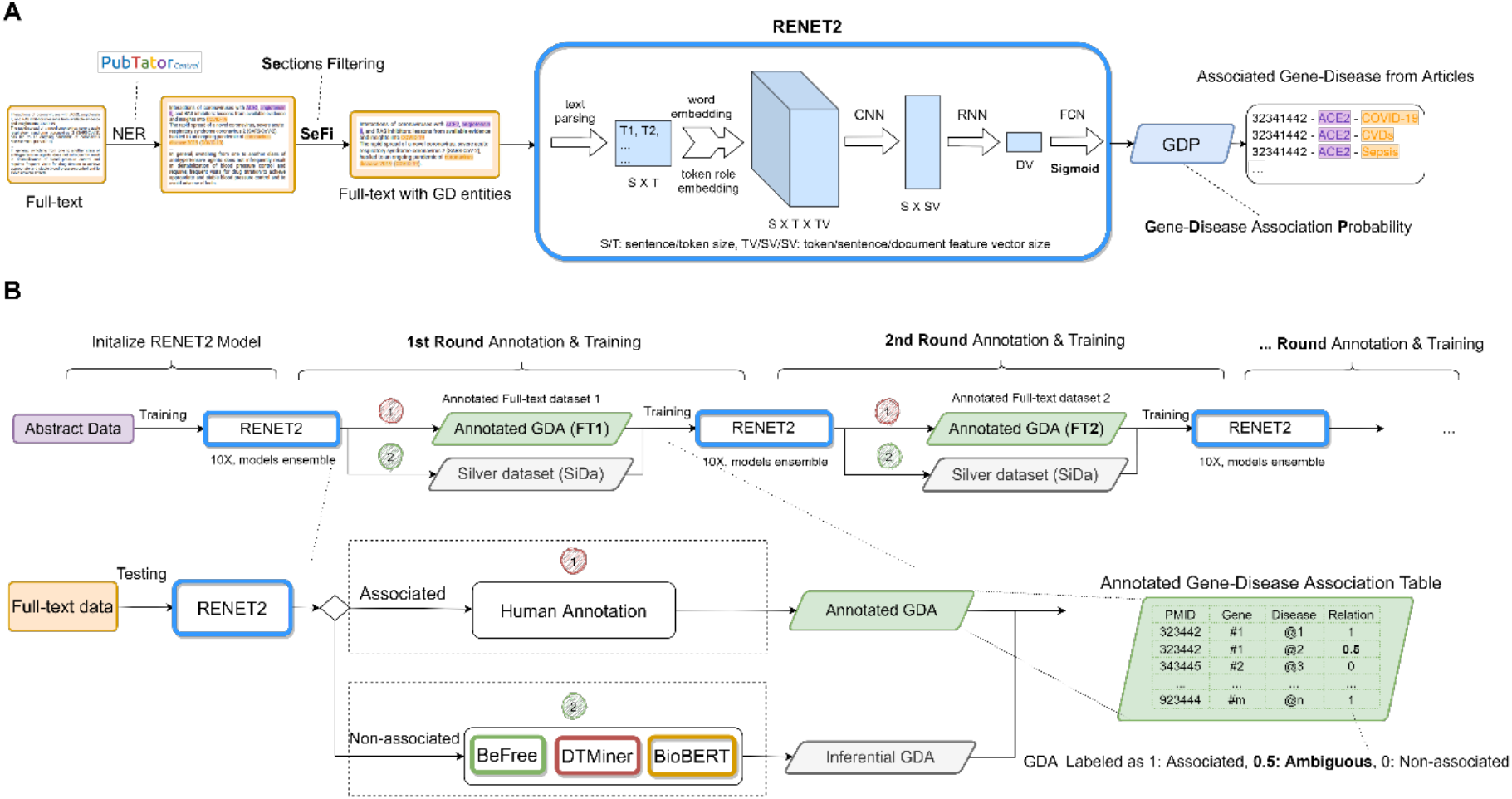
Overview of the RENET2 pipeline. A. Full-text level relation extraction. B. Iterative training data expansion with RENET2 and human annotation.

We compared the performance of RENET2 with the best existing methods: BeFree, DTMiner and BioBERT. We found that RENET2 achieved the best F1 score using a full-text dataset. RENET2 achieved an F1 score of 72.13%, which was 27.22%, 30.30%, 29.24% higher than BeFree, DTMiner and BioBERT, respectively. Using RENET2, we analyzed 1,889,558 full-text articles from PMC and extracted 3,717,569 gene-disease-article associations, which is more than five times the number of associations extracted by RENET from abstract data [14]. To help medical professionals keep track of advances in COVID-19 research [16, 17], we applied RENET2 to the LitCovid article sets, a collection of up-to-date publications on COVID-19. We found 1,231 proteins associated with the COVID-19 disease and ranked the top 15 proteins according to the number of supporting studies. The source code of RENET2, the extracted gene-disease associations, and the proteins associated with COVID-19 with corresponding articles are available at https://github.com/sujunhao/RENET2.

## Materials and Methods

### Dataset descriptions

We used two dataset levels in this study, one at abstract level and one at full-text level. See Supplementary Table S1 for the data on the annotated gene-disease associations. The abstract level dataset comprises 1,000 annotated abstracts and was used 1) as the initial model in iterative training data expansion for training a full-text RENET2 model, and 2) for benchmarking the effect of adding Ambiguous associations. To construct the abstract level dataset, we started with manually annotating all gene-disease pairs in 500 abstracts from scratch. The 500 abstract articles were randomly selected from RENET’s dataset. The annotation process was conducted by three experts with a Biology, Bioinformatics, or Computer Science background, respectively. The workload was distributed evenly. For quality control, an expert can mark an annotation as uncertain. All uncertain annotations were then discussed with and arbitrated by the other two experts. These 500 abstracts should have the highest annotation quality. Using these 500 abstracts, we trained the first RENET2 model and conducted one iteration of training data expansion for another 500 abstracts. For abstracts, we used DisGeNET [18] to assist manual curation. DisGeNET is a public collection of gene-disease associations in abstracts collected from multiple sources but without a unified curation criterion. In the extra 500 abstracts, we regarded the associations found by RENET2 that are also in DisGeNET as true associations, and we manually checked and arbitrated the contradicting associations. The extra 500 abstracts were used only for training, while the fully manually annotated 500 abstracts were used for both training and validation.

The full-text level dataset was constructed starting from the model trained on the 1,000 abstracts to annotate 500 unlabeled full-text articles. The 500 full-text articles are randomly sampled from PMC open access subset [2]. The associated predictions were manually curated, uncertain annotations were dealt the same way as the abstracts, while the non-associated predictions were curated with unanimous support from three other methods: BeFree, DTMiner and BioBERT (see Materials and Methods – Iterative Training Data Expansion for more details). Unlike the procedure used in the previous paragraph, in which another 500 abstracts were included in the second iteration, we used the same 500 full-text articles for data expansion of full-text training data expansion because we found a large space for annotation improvement on the same 500 full-text articles. The number of annotations almost doubled in the second iteration on the same 500 full-text articles (Supplementary Table S1).

## Definition of the problem

The input of the relation extraction problem is article *X*, consisting of *s* tokens *x*_1_, *x*_2_, …, *x*_*s*_. Let *G* = {*g*_1_, *g*_2_, …, *g*_*n*_} denote the set of gene entities and *D* = {*d*_1_, *d*_2_, …, *d*_*m*_} denote the set of disease entities in an article. The task of gene-disease relation extraction is, for gene-disease pair *g*_*i*_ ∈ *G, d*_*j*_ ∈ *D*, to determine whether the article supports an association relation between *g*_*i*_ and *d*_*j*_. The task predicts a relation *y*(*g*_*i*_, *d*_*j*_) ∈ {0, 1}, where 1 represents an Associated relation, and 0 represents a Non-associated relation.

### Ambiguous association

To better model the relationship between genes and diseases, and to represent relationships that are difficult for an expert to decide (because they may be semantically Ambiguous or require extra time for a decision), we introduced an additional relation type, called Ambiguous association, which is defined as a weak or uncertain association between a gene and a disease. As shown in the examples in Figure 2, Ambiguous association lies between an Associated relation (a clear statement or evidence supporting an association) and a Non-associated relation (no statement or evidence supporting an association). Instead of representing an Ambiguous association as a new class, we use a probability score, GDP (Gene-Disease Association Probability). *p*(*g*_*i*_, *d*_*j*_) ∈ [0, 1] to model the new relations. “Non-associated” and “Associated” are still represented by 0 and 1. We use 0.5 to represent an Ambiguous association, as it is between Non-associated and Associated.

**Figure 2.**
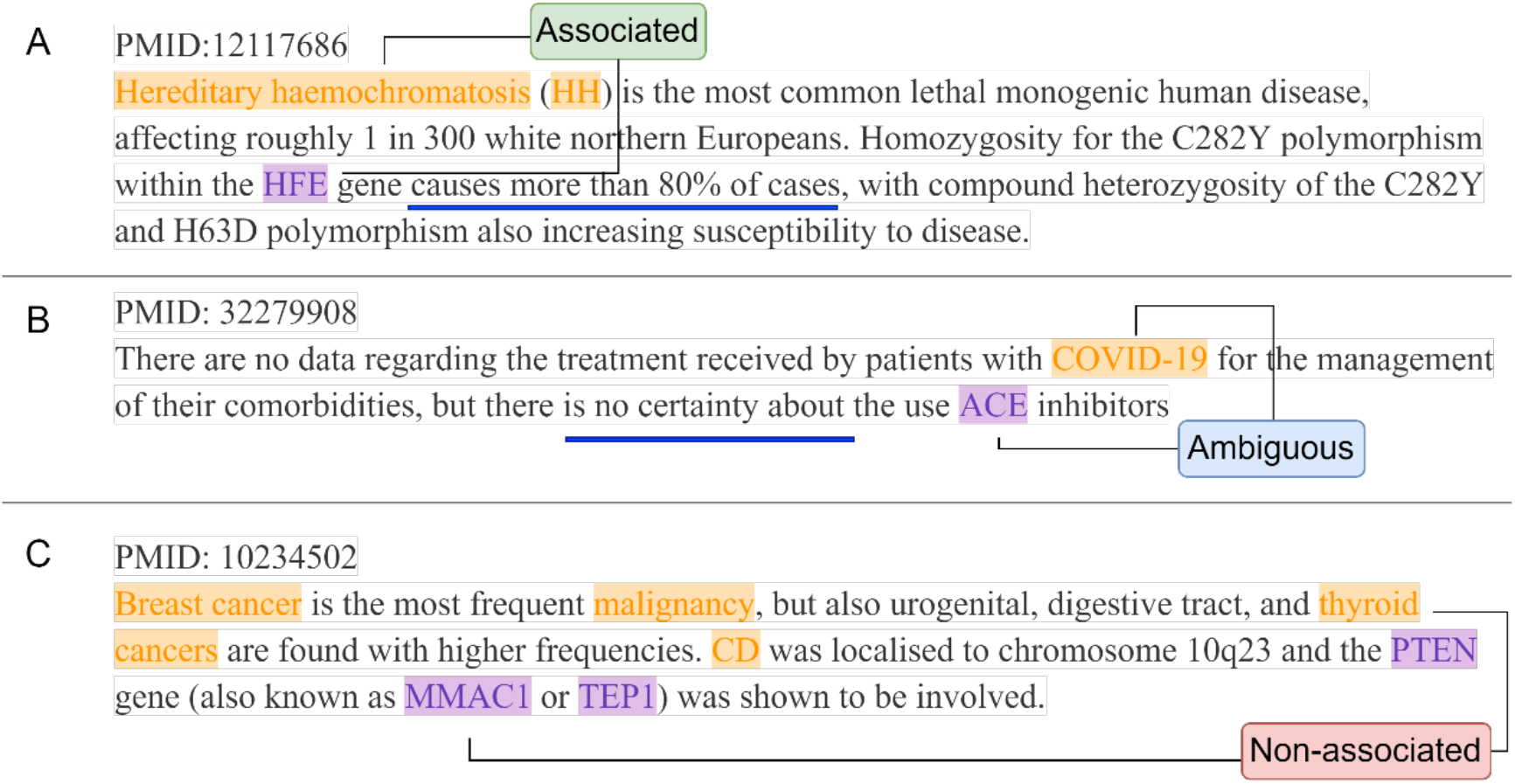
Illustration of the three gene-disease relation types in our dataset. A. Associated relation between *HFE* (gene) and hereditary haemochromatosis (disease) from PMID 12117686 [19]. B. Ambiguous association between *ACE* (gene) and COVID-19 (disease) from PMID 32279908 [20]. C. Non-association relation between *MMAC1* (gene) and thyroid cancer (disease) from PMID 10234502 [21]. Gene entities are purple, and disease entities are yellow.

Ambiguous associations were used in the model training stage to improve the model’s generalization capability. In the prediction stage, we extract Associated gene-disease pairs by computing *ŷ*(*g*_*i*_, *d*_*j*_) as:

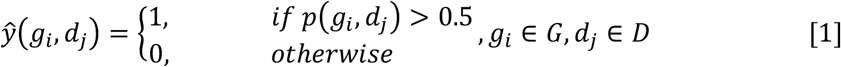

### Overview of the RENET2 pipeline

An overview of the RENET2 pipeline is shown in Figure 1A. First, we conducted NER to identify the gene (*G*) and disease (*D*) entities in an article. For this task, we utilized PubTator Central [22], a state-of-the-art automated concept annotation tool, designed for biomedical full-text articles. It applied cutting-edge machine learning and deep learning techniques for concept disambiguation to improve NER accuracy.

Based on the NER results, we applied section filtering to reduce the input article’s noise. After the preprocessing steps, the data was fed to the RENET2 model for training and prediction. RENET2 produced a probability score, the GDP score, for each gene-disease pair in the article. The GDP scores allowed us to extract the gene-disease associations from the articles.

### Section Filtering (SeFi)

Section Filtering (SeFi) is a technique designed to filter noisy content from full text for relation extraction (RE). We observed that paragraphs containing no gene-disease pairs (i.e. containing only one type or no gene/disease entities), did not provide information on gene-disease relations mentioned at the full-text level. For example, method sections that discussed experiment settings but not specific genes or diseases were not helpful for identifying any gene-disease associations. Based on this observation, we designed a simple filtering technique, called Section Filtering (SeFi), to improve data quality for full-text level relation extraction. The idea of SeFi is to delete paragraphs in each section without any gene and disease entity pair information. It is regarded as a preprocessing module for RENET2. To conduct SeFi, (1) we found all gene and disease entities and paragraph information using PubTator Central, and (2) deleted paragraphs that did not have any gene and disease entity pairs in each section. We found that this technique improved relation extraction performance in full text. See the Results section.

### RENET2 model

RENET2 was built and expanded using the RENET framework. Each word in the RENET2 model is represented by a word vector combined with a one-hot feature vector. The word vector captures the semantic features of a word, while the feature vector denotes whether a word is a target gene, target disease, non-target gene, or non-target disease. Then a document-level representation of the target gene and disease is computed in two steps: (1) from word representation to sentence representation, using a Convolutional Neural Network (CNN), and (2) from sentence representation to document representation, using a Recurrent Neural Network (RNN). Finally, a Feed Forward Neural Network is applied to calculate the Gene-Disease Association Score. For the detailed network architecture and hyperparameters, see Supplementary Note. The computation of RENET2 neural networks is represented as:

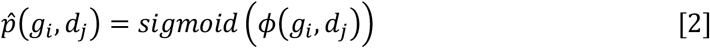

where *ϕ*(*g*_*i*_, *d*_*j*_) is the learned document representation for (*g*_*i*_, *d*_*j*_). Note that we use a sigmoid activation function to compute the probability of association.

To incorporate Ambiguous associations for training, RENET2 models treat extraction as a regression problem. RENET2 uses the mean square errors (MSE) function as the loss function:

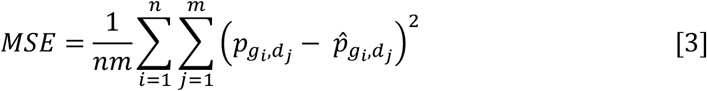

where *m, n* is the number of gene and disease entities in the article, respectively.

We implemented RENTE2 using Pytorch [23]. We estimated that the average token number in a sentence from a biomedical full-text article to be about 27, which is consistent with Lippincott et al. [24]. We configured the maximum token number in a sentence to be 54 to cover most cases. We configured the maximum number of input sentences as 1,000 in a full text. The number was empirically determined, making RENET2 capable of handling a maximum of 54,000 (54 × 1000) tokens, which is more than the number of tokens in most full-text articles. More settings and hyperparameters are available in the Supplementary Note.

### Model ensemble

RENET2 uses the ensemble technique [25] to boost its performance in full-text relation extraction. The ensemble of RENET2 models is done by training *θ* ∈ N^+^ RENET2 models and integrating their prediction results. A gene-disease pair is predicted as Associated if ≥50% of the RENET2 models predict it as Associated. For ensembling *θ* RENET2 models, the ensembled relation type *ŷ*_*ensemble*_ of a gene and disease pair is computed as:

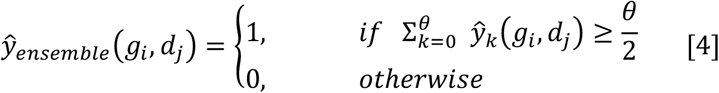

where *θ* ∈ ℕ^+^, and *ŷ*_*k*_(*g*_*i*_, *d*_*j*_) is the relation type prediction of the *k*^45^ model. We set *θ* = 10 as default in RENET2.

### Iterative training data expansion

RENET2, like all other deep-learning methods, requires ample high-quality training samples for good performance. But because of the large number of gene-disease pairs and long text length, manual labeling of full-text articles is labor-intensive and often far too expensive. In RENET2, we developed “iterative training data expansion” to make full-text labeling faster. Instead of manually labeling every full-text article from scratch, the basic idea of the method is to have an expert curate a small number of articles initially labeled by a machine model, and then use the curated labels to train a larger machine model to be used in the next iteration. With the improved performance of the machine model, less and less curation effort is expected to be required in subsequent rounds. The slowdown or even reverse of the machine model performance would denote a stop in curating more machine-labeled results for training. In the field of image classification, a similar idea was used to construct LSUN [26], a large-scale image-classification dataset.

The workflow of iterative training data expansion is depicted in Figure 1B. At the beginning of each iteration, we use the best existing RENET2 model to predict the gene-disease associations from a number of full-text articles. The initial RENET2 model can be from training on a curated public dataset such as DisGeNET, which contains only abstracts, or from the last iteration. Based on the prediction results, all gene-disease pairs are processed using the following two steps:

1. The gene-disease pairs predicted as Associated undergo a manual annotation process. By manually checking each predicted Association, we determine whether it is an Association, Ambiguous association, or Non-association. This step ensures the precision of the positive predictions to be fed to the next iteration. We found this curation process much more efficient than manual-labelling from scratch because 1) most false positive labels are significant to humans, and 2) for labels that trigger hesitation, we simply label them as Ambiguous associations. The annotated gene-disease pairs are regarded as a gold dataset and are used for both training and evaluation.
2. The gene-disease pairs that are predicted as Non-associated are cross-validated by other methods, in our case, BeFree, DTMiner, and BioBERT. We regard a gene-disease pair as inferential Non-associated if all methods predict it as Non-associated. This step ensures the precision of the negative predictions to be fed to the next iteration. As these pairs are cross validated by other methods, but not manually inspected, they constitute a silver dataset (SiDa).

We then enter the next iteration and train a new RENET2 model using both the gold and silver datasets. Depending on the available resources and goal, iteration can be stopped if there is a time constraint or satisfactory performance is reached.

### Evaluation metrics

We use precision, recall and F1 score metrics to evaluate the model performance at each training iteration. Recall can be simply calculated as (annotated associations being correctly predicted) / (total annotated associations). However, calculating precision is more complicated, because we cannot assume all gene-disease pairs in our full-text dataset are annotated, so some true-positive may be misclassified as false-positive. To avoid these misclassifications, we regard an associated gene-disease pair prediction as true-positive if it 1) also exists in our validation dataset and with matched prediction, or 2) is predicted as associated by all four methods: RENET2, BeFree, DTMiner, and BioBERT. Note that we are not saying that a gene-disease association predicted positive by all four state-of-the-art methods is absolutely correct, but we consider that 1) there is a relatively small chance of an association predicted positive by four fundamentally different methods being incorrect, and 2) an association categorized as false-positive in all four methods will have the same effect on changing the precision. The same evaluation methods and metrics were applied to benchmarking all tools. Each experiment was repeated five times with randomly picked 80% training and 20% validation data (i.e. five-fold cross validation).

## Results

### Performance using Ambiguous associations

We expected that if the use of Ambiguous associations in training could improve the performance of relational extraction, it must hold true with abstracts because they usually have the highest density of gene and disease entities. We benchmarked the use of Ambiguous associations with the abstract dataset, and we found it effective in improving both precision and recall. The results are shown in Supplementary Table S2. RENET2 achieved a 2.40% higher F1 score (71.55% against 69.15%) when Ambiguous associations were used. Compared with RENET2’s predecessor RENET, which works only with abstracts, RENET2 achieved a 2.77% higher F1 score (71.55% against 68.78%).

### Performance of iterative training data expansion

We compared the performance of RENET2 at different iterations to evaluate the effect of iterative training data expansion. We trained models using the annotations and curations obtained at the end of each iteration and tested them against our final full-text dataset, i.e., the 500 full-texts with two rounds of curation. The results, shown in Table 1, verified the effectiveness of the iterative training data expansion strategy. With less and less human effort put into each iteration, both precision and recall continued to improve. A leap was observed on recall (from 20.02% to 72.04%) when we switched from using abstracts to full-texts for training, suggesting a substantial difference between abstracts and full texts for extracting gene-disease associations, and the essentiality of having a method designed for full-text RE. From our observation, full texts have, on average, 17 times more tokens and 60 times more gene-disease pairs than abstracts.

**Table 1.** Comparison of RENET2’s performance at different training data expansion iterations. The best result in each column is in bold.

| Training dataset | Testing dataset | Precision | Recall | F1 score |
| --- | --- | --- | --- | --- |
| 500 abstracts | 500 full texts<br>2<sup>nd</sup> curation<br>(5-fold cross<br>validation, 80%<br>training: 20%<br>validation) | 0.6024 | 0.2539 | 0.3573 |
| 1000 abstracts |  | 0.6505 | 0.2002 | 0.3062 |
| 500 full texts<br>1<sup>st</sup> curation |  | 0.6765 | 0.7204 | 0.6977 |
| 500 full texts<br>2<sup>nd</sup> curation |  | **0.7062** | **0.7371** | **0.7213** |

### Comparison of RENET2 models with different training settings

We introduced three new techniques to RENET2: Model Ensemble (ENS), Section Filtering (SeFi) and Silver Dataset (SiDa). But how these techniques work alone and together remained to be studied, so we compared RENET2’s performance with different technique combinations using the full-text dataset. The results are shown in Table 2. We observed that all techniques improved model performance when used alone. SeFi resulted in the most significant improvement in RENET2, since all F1 scores when SeFi was not used were lower than those using SeFi. Using SeFi alone resulted in a 6.57% increase in the F1 score, indicating that filtering out noisy text is critical for full-text RE. In addition, RENET2 excludes the method section for training by default. The influence of each section is analyzed in detail in a later section. ENS alone improved F1 score by 4.05%. SiDa improved F1 score by 1.41%, but it had a different impact on precision and recall, increasing precision by 4.68%, but decreasing recall by 3.25%. This matched our expectation that the additional Non-associated labels from the silver dataset would reduce false positive predictions, but increase false negative predictions to a certain extent. Thus, if SiDa is used for better precision, other techniques are needed to compensate for decreasing recall. When all three techniques were used, the F1 score improved by 11.49% (16.45% higher precision and 4.84% higher recall). Therefore, we will use all three technique in RENET2 by default in future analyses.

**Table 2.**
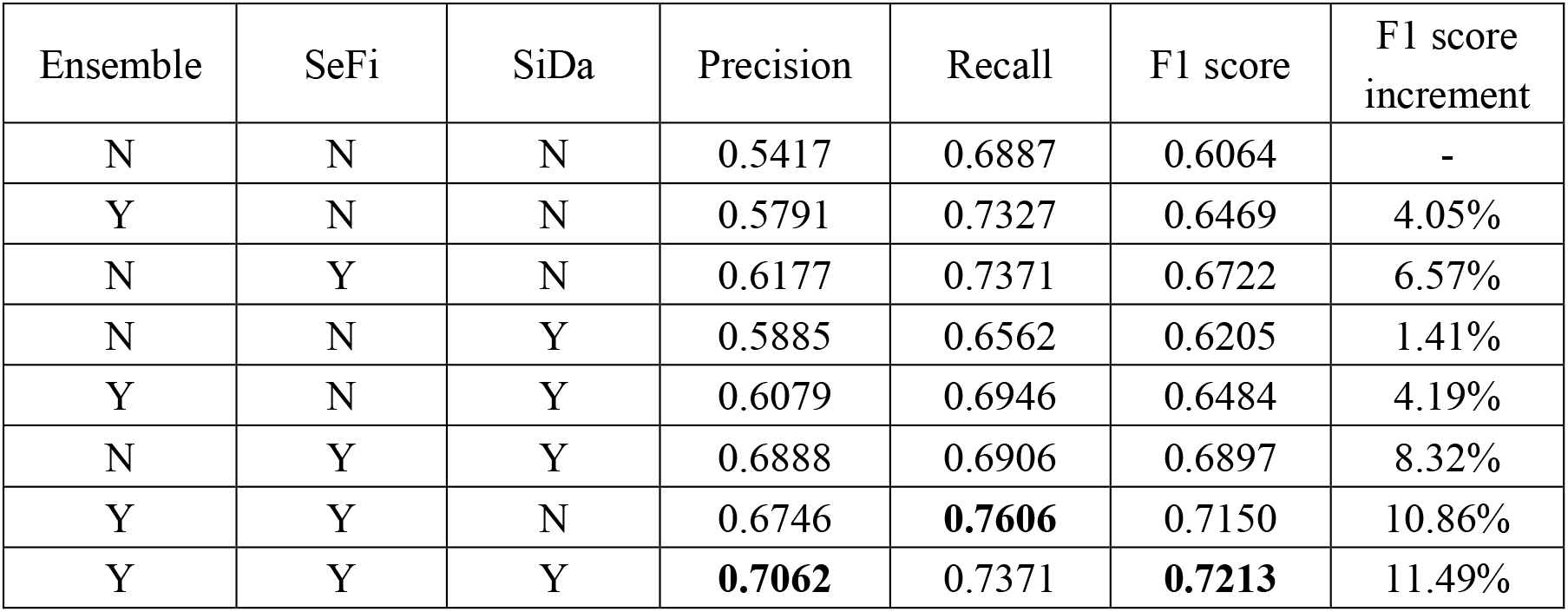
Comparison of RENET2’s performance on the full-text dataset when trained with different settings. The best result in each column is in bold. ENS: Ensemble, SeFi: Section Filtering, SiDa: Silver Dataset, F1 increment: increase in F1 score compared to the basic setting.

| Ensemble | SeFi | SiDa | Precision | Recall | F1 score | F1 score increment |
| --- | --- | --- | --- | --- | --- | --- |
| N | N | N | 0.5417 | 0.6887 | 0.6064 | - |
| Y | N | N | 0.5791 | 0.7327 | 0.6469 | 4.05% |
| N | Y | N | 0.6177 | 0.7371 | 0.6722 | 6.57% |
| N | N | Y | 0.5885 | 0.6562 | 0.6205 | 1.41% |
| Y | N | Y | 0.6079 | 0.6946 | 0.6484 | 4.19% |
| N | Y | Y | 0.6888 | 0.6906 | 0.6897 | 8.32% |
| Y | Y | N | 0.6746 | **0.7606** | 0.7150 | 10.86% |
| Y | Y | Y | **0.7062** | 0.7371 | **0.7213** | 11.49% |

### Comparison between RENET2 and other methods

To the best of our knowledge, RENET2 is the first open-source method optimized for full-text gene-disease RE. We compared RENET2 to three state-of-the-art gene-disease relation extraction methods: BeFree, DT-Miner, and BioBERT. For BeFree and DTMiner, the best pretrained models were downloaded and used for benchmarking. For BioBERT, we used a BioBERT-based model fine-tuned on the GAD [7] dataset. To ensure a fair comparison, all methods used PubTator Central (PTC) for the NER steps.

The results are shown in Table 3. RENET2 outperformed the other three methods by a significant margin, achieving 70.62% precision, which was 39.09%, 43.01% and 42.59% higher than BeFree, DTMiner and BioBERT, respectively. For overall performance, RENET2 achieved a 72.13% F1 score, which was 27.22%, 30.30% and 29.24% higher than BeFree, DTMiner and BioBERT, respectively. The three methods underperformed on precision, partially because they are sentence-based and could not leverage multi-sentence context to sift out non-conclusive associations.

**Table 3.** Comparison of different methods for full-text gene-disease relation extraction. The best result in each column is in bold.

| Method | Precision | Recall | F1 score |
| --- | --- | --- | --- |
| BeFree | 0.3152 | 0.7808 | 0.4491 |
| DTMiner | 0.2761 | 0.8624 | 0.4183 |
| BioBERT | 0.2803 | 0.9128 | 0.4289 |
| RENET2 | **0.7062** | 0.7371 | **0.7213** |
| RENET2 high-sensitivity mode | 0.5518 | **0.9217** | 0.6903 |

Using the default mode of RENET2, we observed lower recall than the other three methods (73.71% against 78.08%, 86.24% and 91.28%). To accommodate some usage scenarios that favor recall over precision, RENET2 also provides a high-sensitivity mode. Using this mode, we achieved the best recall (92.17%) of all methods, while having a 3.10% lower F1 score than the default RENET2. In the high-sensitivity mode, the use of the silver dataset for training was disabled, and the model ensemble’s voting strategy was modified to “predict positive as long as one out of ten models support it”.

### Studying the importance of difference full-text sections for relation extraction

We studied the importance of different sections for full-text RE. We summarized the count of gene-disease associations in each section using the full-text dataset. The results are shown in Figure 3A. We found that introduction, discussion and abstract were the three most informative sections for gene-disease RE. The findings highlight that using abstract alone is insufficient for gene-disease relation extraction because a large portion of gene-disease associations are from the other sections. We also found that few associations were found in the method section.

**Figure 3.**
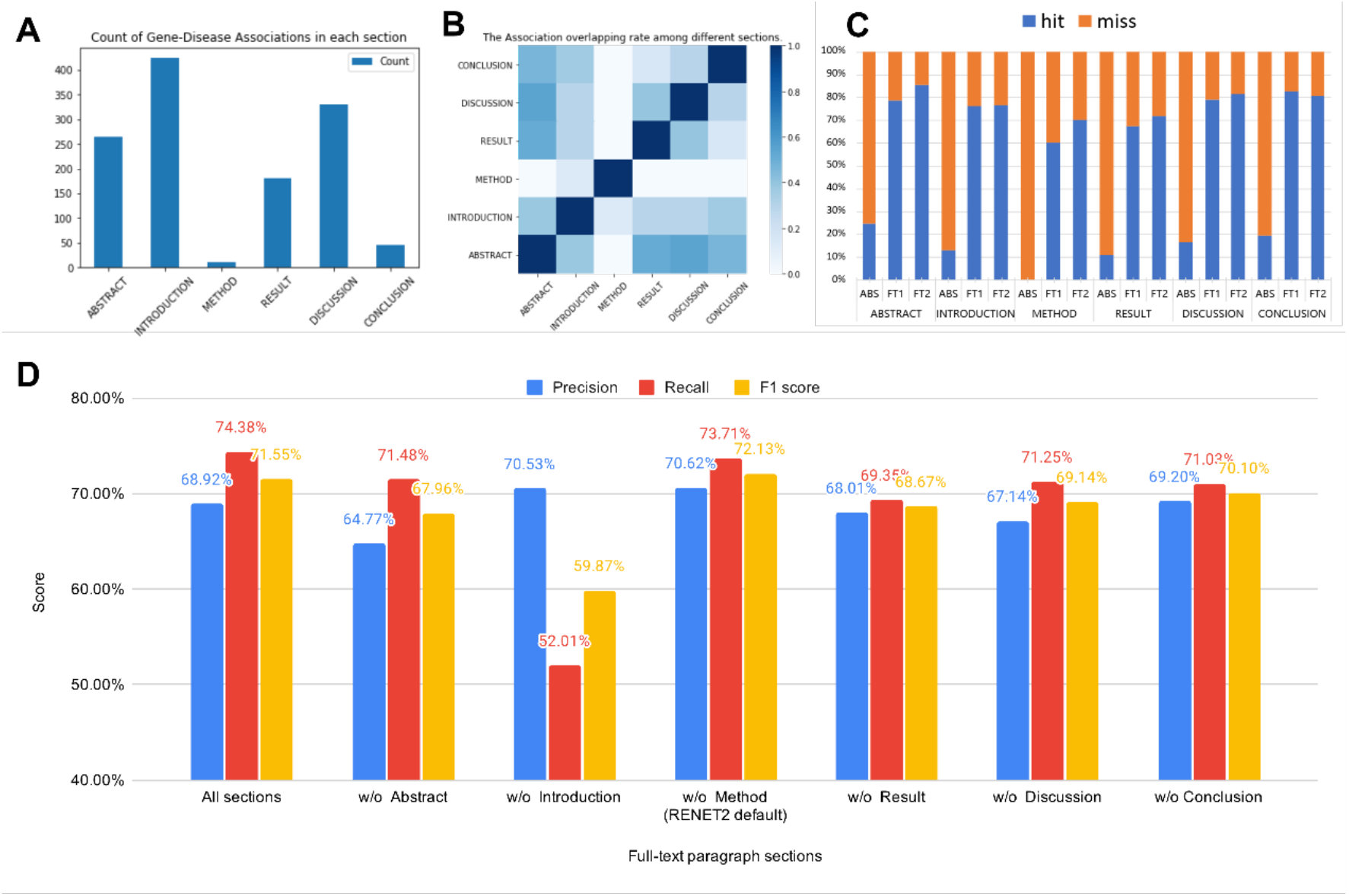
Gene-disease relation extraction in different full-text sections and their effect on RENET2 models. (A) The count of “Associated” predictions in different sections. (B) The association overlapping rate among different sections. The darker color indicates a higher overlap rate. (C) The recall rate of multiple RENET2 models on different sections. ABS: model trained with 1,000 abstracts, FT1: 500 full texts 1st curation, FT2: 500 full texts 2nd curation. (D) RENET2’s performance after removing different sections from training.

To better understand the relationship between the different sections, we measured the overlapping rate of associations found in them. A heat map is shown in Figure 3B. The overlapping rate of two sections is computed as *overlap*(*A, B*) = |*A* ∩ *B*|/ min(|*A*|, |*B*|), where *A* and *B* are the gene-disease associations found in the two sections. We found that the highest overlapping rate was between abstract, result and discussion. This indicates that many gene-disease associations from the abstract can also be cross-validated in the result or discussion section. Although we haven’t made use of this discovery in RENET2, it could lead to even better precision in our future investigations.

To understand how iterative training data expansion improved the recall rate in different sections, we applied three RENET2 models trained with data from different iterations to the full-text dataset. The recall breakdown by section is shown in Figure 3C. We found that the two full-text models performed much better than the abstract model on the abstracts themselves, so we believe that a higher overall number of tokens and a more diverse corpus can comprehensively improve the performance of gene-disease RE.

We also performed an experiment with one-section left out to train RENET2 to see the effect of each section on the prediction power of RENET2. The results are shown in Figure 3D. In spite of removing a section from the training dataset, all models were trained using RENET2’s default setting (i.e. ENS, SeFi, and SiDa enabled). When all sections were used for training, the F1 score was 71.55%. When the method section was left out, the F1 score increased slightly by 0.58%. For this reason, we left out the method section in model training by default in RENET2. We found that leaving out any sections other than method decreased the F1 score from 1.45% to 11.68%. The findings show that leaving out the introduction section severely deteriorated the performance (F1 score from 71.55% to 59.87%), indicating the importance of having the introduction section when training a model for full-text gene-disease RE.

#### Application 1: Large-scale full-text gene-disease relation extraction

We applied RENET2 on a large-scale biomedical literature database to build a collection of gene-disease associations from existing studies. We used RENET2 to extract gene-disease associations from all full-text articles available in the PMC open-access subset (downloaded at 2020/08), which has more than 2.75 million full-text articles. It is the largest collection of full-text articles available for download and text mining [2]. After filtering out articles without any gene-disease pair, we applied RENET2 on the 1,889,558 remaining articles. We found 3,717,569 gene-disease associations from the articles in 14.65 wall-clock hours using a computing cluster with 49 NVIDIA GeForce GTX 1080 Ti GPU cards (detailed statistics in Supplementary Table S3). This was more than five times the number of associations extracted from abstracts by RENET [14]. The scripts for running RENET2 and the entire set of extracted associations are available in RENET2’s GitHub repo.

#### Application 2: Finding proteins associated with COVID-19

With the fast expansion of COVID-19 research, it is getting harder and harder for researchers to keep up with the latest progress. RENET2 provides an efficient way for users to track associated proteins and pinpoint the subset of the literature of interest. We applied RENET2 to extract proteins associated with COVID-19 from the LitCovid [16] dataset. LitCovid is a curated literature hub for tracking up-to-date scientific information about COVID-19. It had 73,654 articles in the dataset dated 2020/06. After filtering out articles without any protein names, we applied RENET2 on the 19,368 remaining articles, and we found 1,231 proteins that are reported to be associated with COVID-19 in at least one article. The results are shown in Figure 4. The top 15 proteins are *ACE2, IL-6, CRP, Spike, ACE, TNF-alpha, TMPRSS2, IL-1beta, ORF1a/b, Fibrinogen, CD8, Mpro, AST, IL-10* and *IFN-gamma*. The findings are consistent with Yeganova et al. [27]. The scripts for running RENET2 and the results are available in RENET2’s GitHub repo.

**Figure 4.**
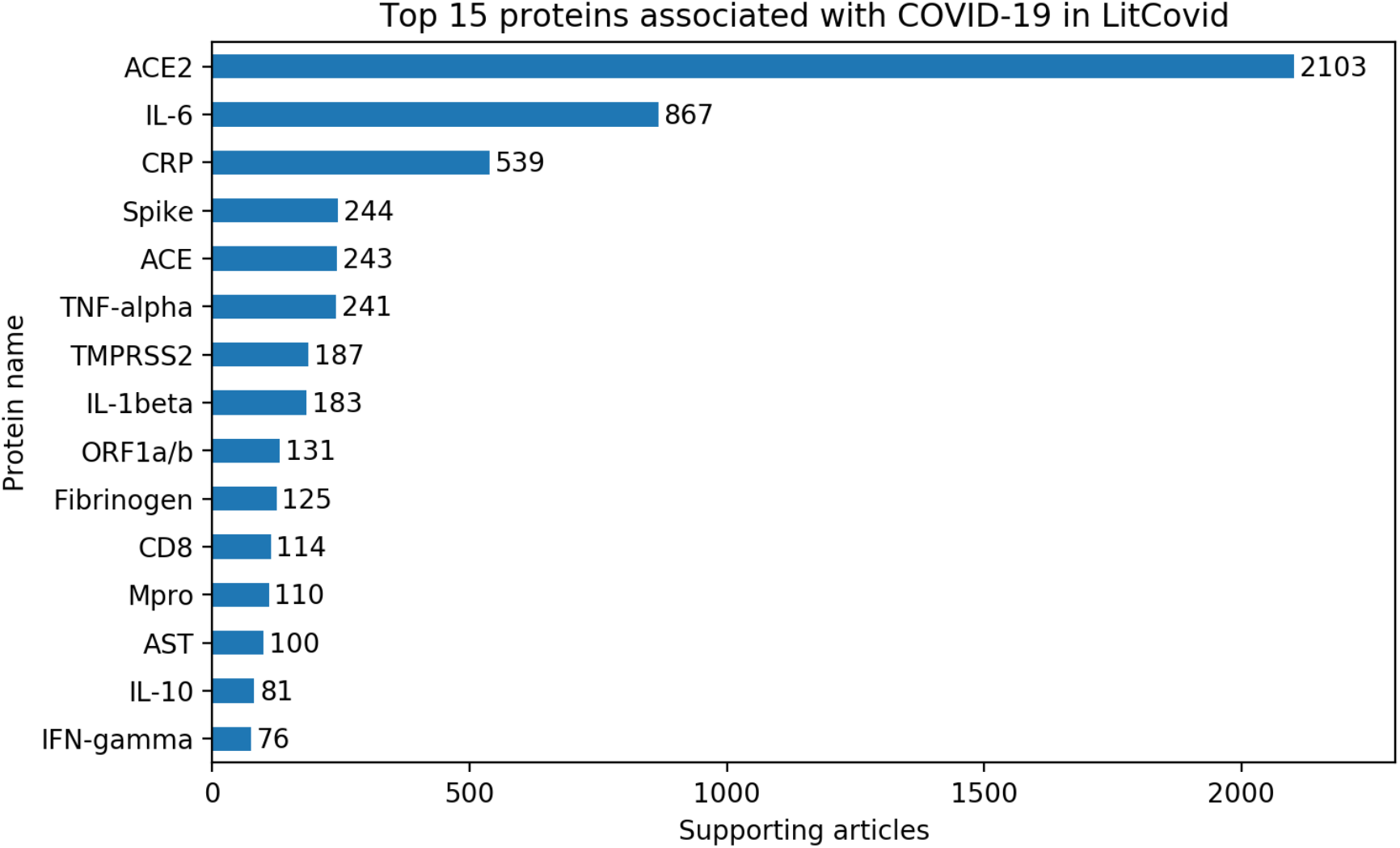
The top 15 genes RENET2 found associated with COVID-19 in the LitCovid dataset. The labels show the number of articles that support the gene’s association with COVID-19.

## Discussion

In this paper, we introduced RENET2, a deep-learning-based relation extraction method to extract full-text gene-disease associations from full-text articles. RENET2 can use Ambiguous associations for training and multiple techniques can be applied, including model ensemble, section filtering and a silver dataset, to boost its performance on full text. The new iterative training data expansion method proved to be effective in improving relation extraction performance, while reducing human effort. In our experiments, RENET2 significantly outperformed state-of-the-art methods on full-text gene-disease relation extraction. We demonstrated RENET2’s utility using two applications. We applied RENET2 to the PMC open-access subset, which includes over a million full-text articles, and extracted over three million gene-disease associations. We applied RENET2 to the fast-expanding pool of COVID-19 research articles and ranked the top 15 proteins verified in another more systematic study. The source code and the results of this study are publicly available in GitHub.

Some practical challenges remain, leaving room for the further development of RENET2. First, the upper-bound accuracy of relation extraction is capped by the accuracy of named entity recognition. Our study used PTC, which is the best system to date for named entity recognition. However, we still found a small number of named entity recognition results to be inconsistent and erroneous, such as incomplete gene and disease entities, and failure to disambiguate gene and disease acronyms. We estimate that the accuracy of the RENET2 model can be improved by at least 8% if the named entity recognition annotation is error-free. Second, the current model is limited to computing one gene-disease pair association at a time, which results in a waste of computation resources [28]. Different gene-disease pairs from the same article share most of their contexts. A significant amount of computing time and resources can be saved if multiple gene-disease pairs are processed in a single computation step. We also hope to incorporate deep language representation models into full-text relation extraction in our future research. Recent studies show that deep language representation models, such as ELMo [29] and BERT [11], can develop strong language understanding capability by pre-training on large-scale unlabeled corpora. We aim to solve the input length limitation, as well as a few other limitations of applying deep language representation models, in the future to improve the performance of full-text gene-disease relation extraction.

## Data Availability

The source-code and the manually curated abstract/full-text training data and results of RENET2 are available at https://github.com/sujunhao/RENET2.

## Acknowledgments

We thank Zhenxian Zheng for his critical comments on model structure and data annotation.

## Supplementary Materials

*Supplementary_materials*.*docx*

