## Supplementary Materials for "RENET2: High-Performance Full-text Gene-Disease Relation Extraction with Iterative Training Data Expansion"

### Supplementary Tables

Table S1. Statistics of (a) the abstract datasets (b) the full-text datasets.

(a)

| Dataset | Documents | Annotated label | | | |
|---|---|---|---|---|---|
| | | Total # | Associated | Ambiguous | Non-associated |
| Abstract | 500 | 2,813 | 870 | 92 | 1,851 |
| Abstract expansion | 500 | 624 | 319 | 47 | 258 |
| Total | 1000 | 3,437 | 1,189 | 139 | 2,109 |

(b)

| Dataset | Documents | Iteration | Annotated label | | | |
|---|---|---|---|---|---|---|
| | | | Total # | Associated | Ambiguous | Non-associated |
| Full-text | 500 | 1st | 793 | 473 | 13 | 307 |
| | | 2nd | 763 | 421 | 3 | 339 |
| | | Total | 1,556 | 894 | 16 | 646 |

Table S2. Comparison of RENET2 for relation extraction from abstracts.

| Model | Using ambiguous associations | Precision | Recall | F1 score |
|---|---|---|---|---|
| RENET | No | 0.6677 | 0.7092 | 0.6878 |
| RENET2 | No | 0.7117 | 0.6724 | 0.6915 |
| | Yes | **0.7127** | **0.7184** | **0.7155** |

The best result of each column is in bold.

Table S3. Data and computational resource consumption statistics of RE from 1,889,558 PMC open-access full-text articles.

| **Data** | |
|---|---|
| Number of Full-text | 1,889,558 |

| | |
|---|---|
| Number of Found GDAs | 3,717,569 |
| Raw Data Size | 15.4 GB |
| **Computational resourse consumption** | |
| Total wall-clock | 14.65 hours |
| Total GPU hours | 718 hours |
| GPU Cards (NVIDIA GeForce GTX 1080 Ti) | 49 |

### Supplementary notes

### The RENET2 network architecture

RENET2 in sequence uses an embedding layer, a convolutional neural network (CNN), a recurrent neural network (RNN), and a fully-connected network (FCN) to compute the probability of GDA.

RENET2 uses embedding layers to map token IDs to a vector of features, which consists of two parts: 1) word embedding for each token's context meaning, and 2) role embedding to indicate a token's role in the sentence, including target gene, target disease, non-target gene, and non-target disease. Word embedding and role embedding are concatenated together as the word representation input for the next layer. The input size settings of RENET2 are as follows:

| | Abstract | Full text |
|---|---|---|
| Max sentence | 32 | 1000 (basic), 400 (SeFi) |
| Max tokens per sentence | 54 | 54 |

RENET2 uses CNN to obtain the sentence feature, BiLSTM to obtain the document feature, and three-layer FCNs, followed by a sigmoid activation, to compute the probability of each GDA. The network architecture layer settings are as follows:

| Layers | Size |
|---|---|
| Token ID embedding | 64 |
| Token role embedding | 4 |
| CNN kernel size (Conv1D) | [2, 3, 4, 5] X 100 |
| RNN hidden size (BiLSTM) | 69 X 2 |
| FCN1 | 136 X 136 |
| FCN2 | 136 X 136 |
| FCN3 | 136 X 1 |

The hyperparameters used in RENET2 for different types of inputs are as follows:

| Hyperparameters | Abstract | Full text |
|---|---|---|
| Learning rate | 1e-3 | 8e-4 |
| L2 penalty term | 1e-4 | 5e-5 |

| | | |
|---|---|---|
| Training epochs | 18 | 10 |
| Dropout rate, after token ID embedding | 0.3 | 0.3 |
| Dropout rate, after FCN2 | 0.1 | 0.1 |
